# Circulatory systems and mortality rates

**DOI:** 10.1101/2021.05.27.446029

**Authors:** Gurdip Uppal, Pinar Zorlutuna, Dervis Can Vural

## Abstract

Aging is a complex process involving multiple factors and subcellular processes, ultimately leading to the death of an organism. The microscopic processes that cause aging are relatively well understood and effective macroscopic theories help explain the universality of aging in complex systems. However, these theories fail to explain the diversity of aging observed for various lifeforms. As such, more complete “mesoscopic” theories of aging are needed, combining the biophysical details of microscopic failure and the macroscopic structure of complex systems. Here we explore two models: (1) a network theoretic model, and (2) a convection diffusion model emphasizing the biophysical details of communicated signals. The first model allows us to explore the effects of connectivity structures on aging. In our second model, cells interact through cooperative and antagonistic factors. We find by varying the ratio at which these factors affect cell death, as well as the reaction kinetics, diffusive and flow parameters, we obtain a wide diversity of mortality curves. As such, the connectivity structures as well as the biophysical details of how various factors are transported in an organism may explain the diversity of aging observed across different lifeforms.

## INTRODUCTION

In many complex organisms, including humans, the probability of death (mortality rate) *µ*(*t*) typically increases exponentially with age *t*. This empirical trend, *µ*(*t*) ∼ *e*^*λt*^ is known as the Gompertz law [1]. However, a broader look across taxa reveals mortality curves that are as diverse as life itself [2]. Most interestingly, there appears to be phylogenetic correlations in the shape of *µ*(*t*) [3]: Mammals tend to have steep mortality curves that increase many folds during their lifespan, which means that an old mammal is many times more probable to die compared to a mammal of average age. In contrast, for amphibians and reptiles *µ*(*t*) changes little within their lifetime. Plants tend to age even less, and some can even anti-age, i.e. exhibit mortality rates that decrease with age. Invertebrates on the other hand, are scattered across the map: Some, such as water fleas and bdelloid rotifers age as rapidly as mammals, while others, such as the hydra and hermit crab do not seem to age. What is the reason behind these phylogenetic trends? What taxa-specific anatomical features set the characteristics of the mortality curve of an organism?

Evolutionary theories explain aging as due to selection failing to remove deleterious mutations whose effects manifest after peak fertility (mutation accumulation theory) [4, 5] or due to mutations that increase fitness and reproductive potential early in life at the cost of that later in life (antagonistic pleiotropy theory) [6–8]. These standard theories predict that mortality rate should rapidly increase after maturation, and therefore cannot account for the variations in mortality curves, nor the mechanism behind phylogenetic correlations.

On the other end of the spectrum there are mechanistic theories of aging, seeking to understand aging in terms of biochemical basic principles such as telomere shortening [9], cell damage and repair, [10], oxidative stress [11], and cell metabolism [12]. While these factors surely drive aging in all organisms at the cellular level, it is unclear why they manifest differently on the macroscopic scale depending on the bau-plan.

Between these two extremes of evolutionary and biochemical theories, there are “mesoscopic” theories of aging, which attempt to bridge cellular-level failures with organismic failure. Reliability theory [13] views the organism as a collection of clusters (e.g. organs) of smaller units (e.g. cells), and assumes that if all units within any one of the cluster fails, then so does the organism. The network theory of aging [14], views the organism as a large collection of units that depend on one another to function. Once a unit malfunctions, so will its dependents. As such, microscopic failures that otherwise accumulate slowly, can suddenly cascade into a macroscopic catastrophe.

The goal of this paper is to expand the interdependence network theory of aging [14] to develop a new hypothesis that can account for the taxa-specific characteristics of mortality curves. The main idea is that in a biophysically realistic interdependence network, the structure of connections between the nodes are determined by the exchange of various types of goods and signals, as mediated by diffusion and circulatory flow. Therefore, the transport system within an organism (by virtue of being a proxy for the interdependency between functional body components) will govern how it fails. We will argue that the qualitative differences in the mortality curves may be due to (a) the structure of the interdependence network and (b) the type of goods exchanged through the network.

The simplest species rely primarily on diffusion to transport necessary resources into and waste out of the organism. As specialized organs form, complex species evolved circulatory systems to more effectively transport required resources between components in the organism. This allows for stronger interdependencies to form within an organism, leading to stronger aging effects. For example, in a sparse decentralized interdependence network such as that of a bush or ivy, the function of a leaf or terminal branch in one location does not critically depend on another one far away. Because of this decentralized structure, accumulated failures will only influence near-by nodes and not lead to a catastrophic cascade. The bauplan of a mammal on the other hand is a dense, many-to-many interdependence network. Virtually every functional component couples to every other via blood stream, and therefore the failure of few components will rapidly cascade into a catastrophe. Thus, our first specific hypothesis is that organisms with a large number of components that are functionally coupled irrespective of the distance between them, will exhibit steeper mortality curves.

In the first part of our paper we will simulate the failure modes of cohorts of networks that vary in structure, to illustrate that dense, many-to-many networks (as in vertebrates) exhibit steep mortality curves, whereas sparse, few-to-few networks (as in plants) exhibit flatter or even decreasing mortality curves.

Secondly we will argue that the type of products exchanged between nodes make an important difference in mortality curves. For example, in reptiles and amphibians, oxygenated and venous blood mixes, whereas in mammals, it does not. As such, for a mammal, the death of every cell is a small step closer to organismic failure, whereas for a reptile or amphibian, this infinitesimal loss of function may be partly compensated by lesser CO_2_ stress.

Accordingly, in the second part of our paper we will examine the mortality statistics of cell populations that exchange only positive factors versus a mixture of positive and negative factors. We will do so by simulating the demographics of a population of simplified organisms where cells are arranged around a circulating fluid. In one population, clean and dirty blood will not mix, so cells will only be subject to each other’s positive effects (growth factors, survival factors, antioxidant enzymes [18–21]). We will see that this population will exhibit a rapid aging curve, i.e. mortality rate will be many times more larger at average lifespan compared to that at half of lifespan. This population will be compared with a “mixed blood” one, where cells will be subject to each other’s positive as well as negative factors. We will see that this population will exhibit weak aging characteristics, i.e. will have flatter mortality curves.

We find that the biophysical details such as type of factor, reaction kinetics, and diffusion and flow rates, can drastically affect the aging dynamics of a system, even for a schematic circular circulatory system. Our results emphasize the importance of biophysically accurate models in understanding the interdependence structure of an organism and the aging dynamics of complex living systems.

## METHODS

To explore the effects of the interdependency structure on aging dynamics, we study two models of aging. One where the interdependency between body components are governed by a complex network structure, and one where it is described by a simple circulatory flow.

### Network model

In the first model, we simulate aging of interdependency networks that have a variety of structures to study how local and global couplings between components (network nodes) shape the mortality curves of a species. The nodes in a network can represent genes, cells, organs or some collection of components that has a specialized function in an organism; whereas edges between nodes represent dependencies. Each node is assumed to be in either a functional or nonfunctional state. The model is implemented as follows: (1) we create a random directed network. We explore various topologies as shown in Fig. 1. The scale free random networks in Fig. 1d,e, were generated using the Barabasi-Albert model with a given initial number of connected nodes *m*_0_. The random networks in Fig. 1e,f, were generated using the Erdos-Renyi model varying the probability of an edge *p*. The fully connected network in Fig. 1h was generated by attaching one edge between any two nodes, with the direction chosen randomly. We also assign an initial fraction of nodes, *η* to be in a nonfunctional state. (2) We age the network system by first randomly turning functional nodes into non-functional ones with probability *π*_*d*_. We then go through “repair” stage, where nonfunctional nodes turn into functional nodes with probability *π*_*r*_. Next, we cascade the failures: all nodes whose majority of inputs are non-functional turns into a non-functional node. We iterate this procedure of damaging, repairing and cascading till only a small fraction of nodes *τ* remain functional, after which the organism is considered to be dead. Throughout, we set *τ* = 0.01, corresponding to 1% of initial number of nodes. We also set *π*_*d*_ = 0.002, and vary *η* and *π*_*r*_. We study an ensemble of 10000 networks for each topology and record the death times to determine the mortality rate over time (Fig. 1).

**FIG. 1.**
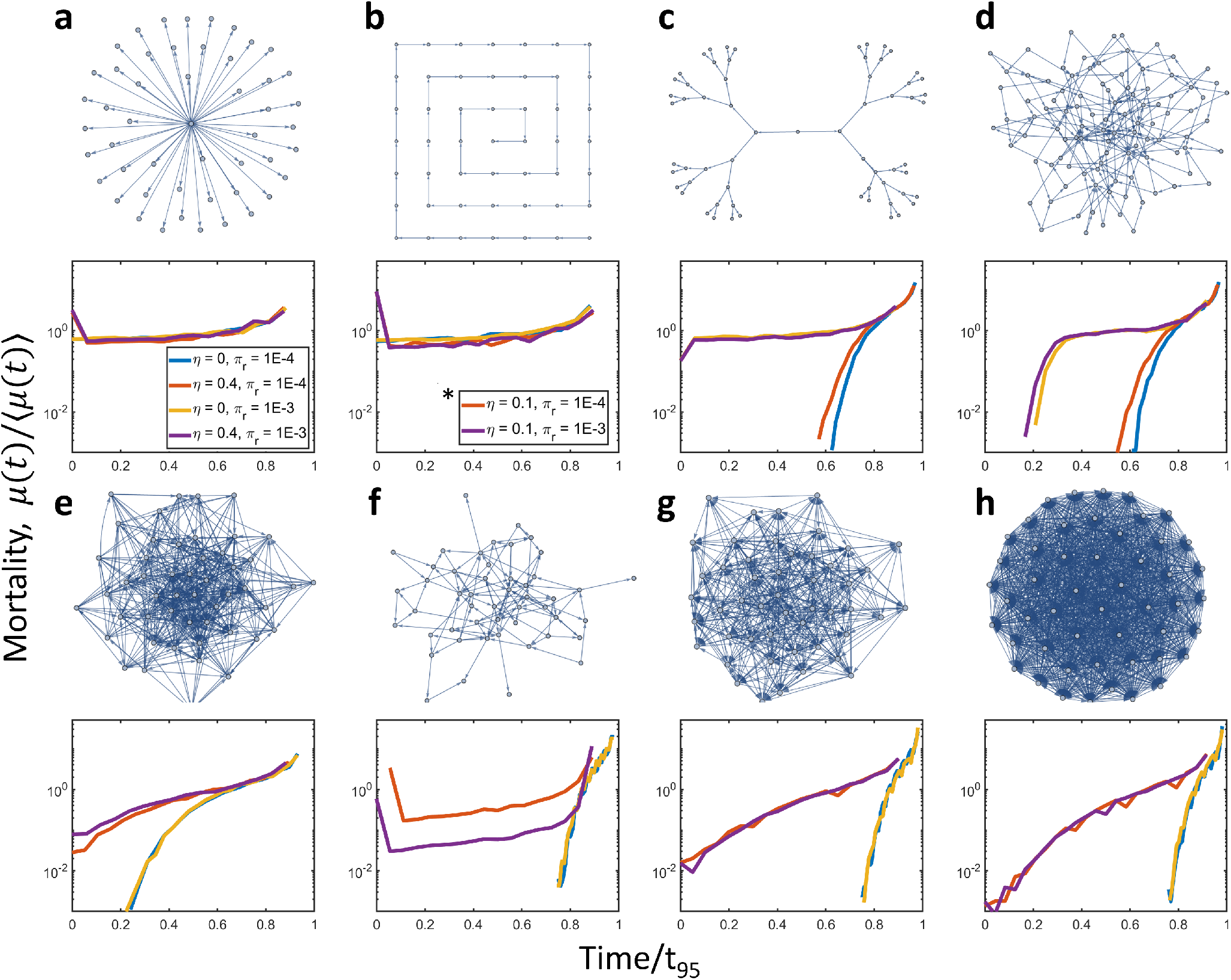
Aging in various network topologies. **a**, Network where one central node is connected to all others. Except for the central node all other nodes are essentially independent. Individual nodes primarily die independently, unless the central node fails, which leads to a system-wide failure. Hence, the mortality curve is mostly flat. **b**, Nodes connected in a chain. With this topology, the failure of a node will cause all nodes “downstream” to fail as well. **c**, Network with branching topology where outer nodes depend on inner ones. The entire network depends on a central node either directly or indirectly. **d**, Scale free random network with low connectivity (*m*_0_ = 2). **e**, Scale free random network with high connectivity (*m*_0_ = 50). **f**, Random network with low connectivity (probability of a link, *p* = 0.1). **g**, Random network with high connectivity (probability of a link, *p* = 0.5) **h**, Fully connected randomly directed network. In general, networks that are less connected have flatter mortality curves, whereas more connected topologies lead to steeper aging curves. The fully connected topology **(h)** gives the steepest aging curves, and the “one-to-all” topology **(a)** gives the flattest curves. In addition to varying topologies, we explored varying initial conditions, by varying the intial fraction of nonfunctional nodes *η*, and varying repair rates *π*_*r*_. We see for more locally connected networks, the repair rate has a much larger effect. In panels **c**,**d**, we see that the repair rate determines the aging curve and the initial condition has little effect. Whereas for more globally connected networks **e-h**, a larger repair rate does little to shape the mortality curve. The initial condition on the other hand, has a larger effect in these cases and less so on locally connected networks. For the simple topologies of **a**,**b**, the mortality curves are predominately flat and nearly independent of parameters. Images of the different networks are shown with limited edges for clarity. For simulations, the branching network was chosen with 1023 nodes, and all other models were chosen with 1000 nodes. An ensemble of 10000 “organisms” was simulated for each case to determine mortality curves. Mortality curves were calculated using 1*/*4 the standard deviation of death times as a time step (*δt*) and taking a discrete derivative of the survival function (*µ*(*t*) ≈−[*S*(*t* + *δt*) −*S*(*t*)]*/*[*S*(*t*)*δt*]) and then normalized by mean mortality rate over time. *The initial fraction of nonfunctional nodes is chosen to be *η* = 0.1 for the red and purple curves in chain topology **(b)**, since much larger *η* leads to nearly immediate failure of the whole network.

### Diffusion-advection model

In the second model, we study the role of the type and transport-mode of signals being communicated, focusing on a simple schematic circular topology. In this model, dependencies are mediated by diffusive and circulatory processes. Cells interact through secreted cooperative factors and/or antagonistic factors. Positive factors help other cells survive or function, whereas antagonistic factors harm or hinder cellular function. These factors then diffuse, flow, and decay within the circulating fluid. We quantify these model assumptions with the following system of equations,

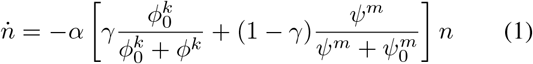

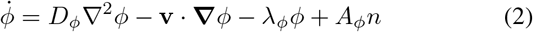

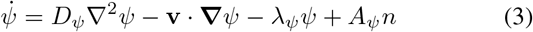

Here *n* = *n*(*x, t*) denotes the cell concentration at time *t* and position *x, ϕ*(*x, t*) is the density of cooperative factors, and *ψ*(*x, t*) is the density for negative factors. Here it is assumed that the cell repair and death are influenced by *ϕ* and *ψ* according to the Michaelis-Mentin (Hill) functions. We hypothesize the ratio of positive versus negative factors mixed in the circulatory fluid will in general vary across species and have a significant effect on the aging dynamics of that species. To study the effects of varying the ratio of positive over negative factors, we introduce a mixing factor *γ* ∈ [0, 1] in equation 1 The constant *α >* 0 gives an overall scaling of the death rate of cells. Note that the two terms on the right hand side of equation 1 could have had two independent coefficients *a, b* instead of *γ* and 1 − *γ*, since setting *α* = *a* + *b* and 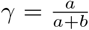 leads to 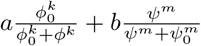.

The constants *ϕ*_0_ and *ψ*_0_ (which we set to 1 throughout) give the saturation constants for positive *ϕ*(*x, t*) and negative *ψ*(*x, t*) factors respectively. These correspond to the “required amount” of factors for a cell to function normally for positive factors, and a “tolerance threshold” for negative ones, above which cells begin to die faster. The constants *k*_*ϕ*_ and *k*_*ψ*_ describe the steepness of the response of cells to the corresponding factor. The constants *D* give the diffusion rate for each chemical (denoted by subscripts), *v* gives the flow rate, taken to be constant and the same for all factors, *λ*’s correspond to decay rates, and *A*’s give the rate at which cells produce the factors.

In our numerical simulations, we treat cells as discrete points that die stochastically with probability given by the term in square brackets in equation 1. The chemical factors are treated as continuous fields that evolve according to a finite difference scheme on a one-dimensional circle with length *L* = 100, following periodic boundary conditions. More specifically, at each time step Δ*t*, cells secrete each factor at a rate given by *A*_*ϕ*_ for positive factors and *A*_*ψ*_ for negative ones. Cells then die with probability given by 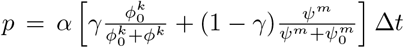. The chemicals dif- fuse, flow, and decay according to a forward Euler finite difference scheme using a central difference for diffusion and a first-order upwind scheme for advection. Lifespan is defined as the time at which all cells are dead. Each system is run multiple times, recording the death time for each to give survival *S*(*t*) and mortality curves *µ*(*t*).

## RESULTS

### Effects of topology and connectivity on aging

We first study the effects of different connectivity structures on the aging of organisms. We explore the effects of different initial conditions and bauplans by varying network topologies and the initial fraction of nonfunctional nodes *η* (Fig. 1). For sparser, locally connected networks, we see the mortality curves are much flatter compared to dense, globally connected networks which have much steeper mortality curves overall (Fig. 1). For densely connected networks, we also see that organisms with poor initial conditions (higher *η*), have flatter mortality curves compared to similarly structured organisms with better initial conditions (lower *η*). This is because these networks are more largely affected by initial conditions, and the probability of failure is already high with poor initial conditions, and does not increase much further with time. For very sparsely connected networks, we also see an initial dip due to higher infant mortality rate from poor initial conditions (Fig. 1a,b). As nodes are repaired, the mortality rate quickly decreases, and then continues to increase as the organism ages. For intermediate connectivities, we see the initial condition does not effect the aging curves (Fig. 1c,d). For denser, globally connected networks, we do not see a large difference from varying the repair rates *π*_*r*_ (Fig. 1e-h). However, for sparser, locally connected networks (Fig. 1c,d), we see higher repair rates lead to flatter mortality curves. This is because failure propagates slower in less connected topologies, and is more amenable to repair.

We therefore find that the connectivity structure, initial conditions, and regenerative capabilities of an organism can greatly influence its aging dynamics. Simpler organisms that are only locally connected age slower or do not age, whereas more complex species with more global connectivity structure have much steeper mortality curves (Fig. 1). What can we say about differences in mortality patterns between species of similar complexity? From the network-theoretic viewpoint, reptiles and amphibians are also complex multicellular organisms, yet we observe them to age much slower than many mammals. How do we then explain the differences in aging across complex organisms, and what are the roles of diffusive and advective processes? Below, we will see that in additional to the topology of interactions, the *type* of interactions and the mode of transport mediating the interactions also play an important role.

### Effects of hill constants and mixing on aging dynamics

To study the effects of the types of signals communicated in an organism, as well as their transport properties and reaction kinetics, we next explore aging patterns of a group of cells situated around a circulating flow, which mediates their interactions.

We start by investigating the effects of the hill parameters and mixing factor *γ* on the aging dynamics of a species. We plot the cell population, total number of living organisms, and mortality rate versus time, normalized by mean lifetime in Fig. We vary both the hill constants and the mixing factor *γ*. For simplicity, we take the two hill parameters to be equal, *k*_*ϕ*_ = *k*_*ψ*_ = *k*. We also set *D*_*ϕ*_ = *D*_*ψ*_ = 5, *α* = 100, and set flow rate to zero. Note that changing the constant *α* amounts to a global rescaling of time. Since we rescale time by the mean lifetime, the constant *α* does not play a major role in our results.

For each parameter set, we run 1000 simulations and record the time at which all cells die as the death time of a species. We plot the total number of species alive (out of 1000) versus time in the middle row of Fig. 2. For the mortality rate, we numerically compute *µ*(*t*) =— [*S*(*t* + *δt*) − *S*(*t*)]*/*[*S*(*t*)*δt*], where we choose the time step *δt* to be given as 1/4 times the standard deviation of the mean lifetime for each species.

**FIG. 2.**
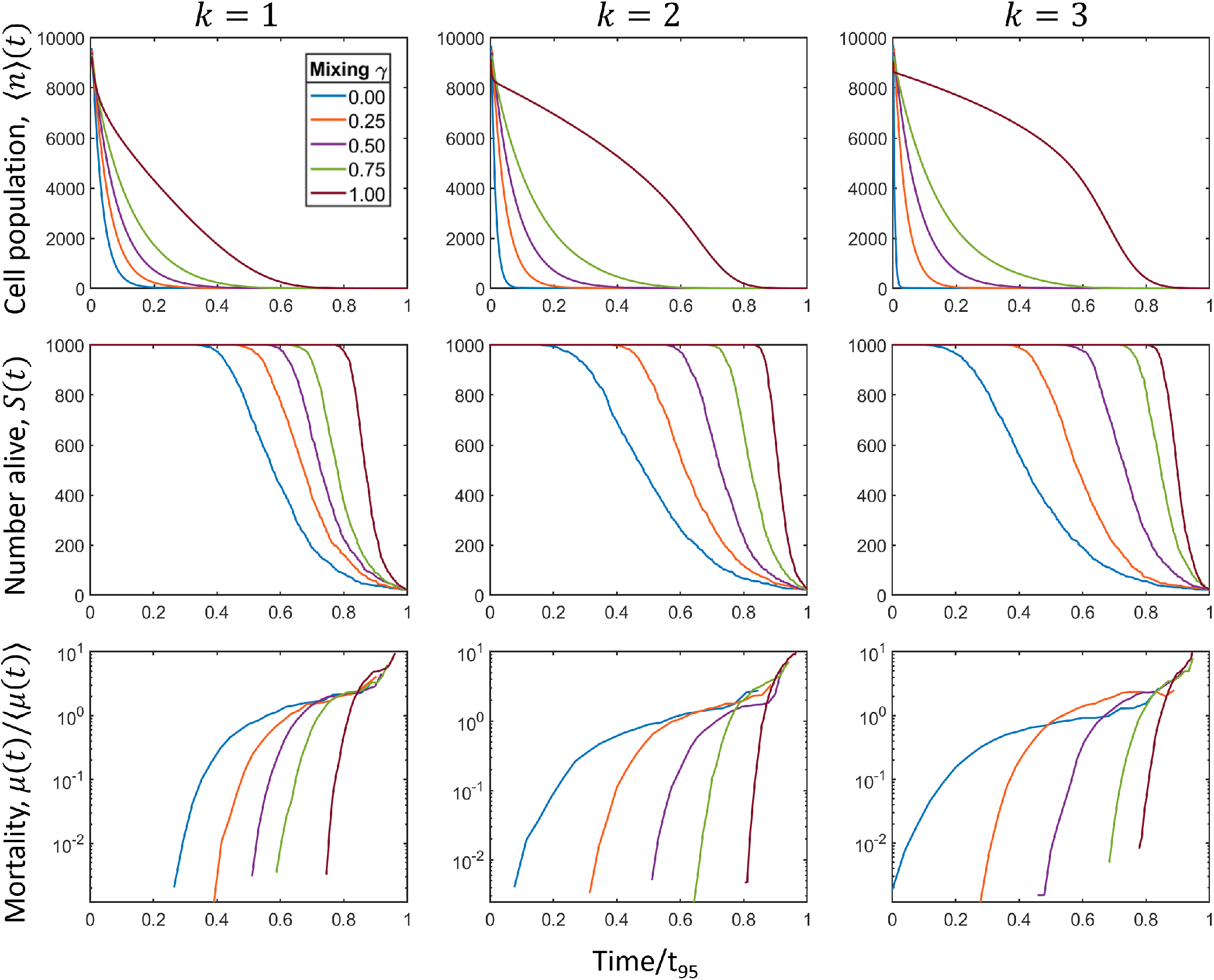
Effects of hill constants and mixing factor *γ* on aging dynamics. We explore the mean cell population over time as well as the aging dynamics for ensembles of simulations, for varying hill constant and mixing factor parameters. For simplicity, we take *k*_*ϕ*_ = *k*_*ψ*_ = *k*. In the top row, we plot the mean cell population over time, normalized by the mean lifetime for each system. For systems where cells communicate more through negative factors (small *γ*), cells die quicker at first and then much slower later. This corresponds to shallower mortality curves, as seen in the last row. For systems where cells communicate more through cooperative factors (large *γ*), cells die out gradually at first and then exhibit more of a crash. This corresponds to steeper mortality curves in the last row. As we increase the hill constant *k*, the effects become more pronounced. For large *k*, cooperative systems age more and antagonistic systems age even less. Survival curves *S*(*t*) in the middle row are given by recording the death times for 1000 simulations for each parameter set. Mortality curves were calculated using 1*/*4 the standard deviation of death times as a time step (*δt*) and taking a discrete derivative of the survival function (*µ*(*t*)≈− [*S*(*t* + *δt*) −*S*(*t*)]*/*[*S*(*t*)*δt*]) and then normalized by mean mortality rate over time. Other parameters were set to *D*_*ϕ*_ = *D*_*ψ*_ = 5, *v* = 0, *α* = 100, *λ*_*ϕ*_ = *λ*_*ψ*_ = 50, and *A*_*ϕ*_ = *A*_*ψ*_ = 10.

We normalize the mortality rates by the mean mortality over time and normalize time by the mean lifetime to compare each case.

The key quantity of this model is *γ*. A *γ* value close to 1 indicates that cells cooperate by secretions that enhance each other’s survival. As we decrease *γ* towards 0.5 this corresponds to cells both benefiting as well as harming each other. Values of *γ* closer to zero describes a collection of cells that solely compete with each other, as one might expect in a colony of non-social unicellular organisms.

As expected, we see that the mortality curves become steeper as we increase *γ*. This is because as we increase *γ*, the interaction between cells go from being deleterious to cooperative. When cells are strongly dependent on each other for survival, local failures can quickly spread to global ones, leading to system wide collapse. For low *γ* on the other hand, cells influence each other negatively. As some cells begin to die, there is less waste pollution and others can survive longer. The mortality curves are therefore more shallow for low *γ*.

The nature of the interaction also manifests in the cell population dynamics. In the top row of Fig. 2, we plot the mean cell population versus time, normalized by mean lifetime. We see for more cooperative systems (higher *γ*), cells die slower at first and then have more of a “crash” near the end of the organism’s lifetime. Whereas for more competitive systems (lower *γ*), cells die quickly at first, and then slowly fizzle out towards the end of the organism’s lifetime.

As we increase the hill constant *k*, we see the same trends with the factor *γ*, but the effect becomes more pronounced. For larger *k*, the interaction between cells increases. Hence, large *γ* systems age even more (steeper mortality curves) and low *γ* systems age even less (shallower mortality curves).

We therefore see that the type of interaction and the reaction kinetics between cells in an organism can strongly influence the mortality of an organism. Even though all organisms are represented by the same topological system in the model, we see that the biophysical details of the interactions also plays a major role in aging.

### Effects of diffusion and flow

We next study the effects of varying diffusion and flow rates on the aging dynamics of organisms. As diffusion and flow rates will alter the extent to which cells interact and couple to each other, we expect the aging dynamics of a species to also vary with different flow and diffusion rates.

For simplicity, we focus on purely cooperative (*γ* = 1) and purely antagonistic (*γ* = 0) interactions. To compare the aging dynamics for varying diffusion and flow rates, we plot the mortality at the mean lifetime of a system versus different diffusion rates and flow rates in Fig. 3.

**FIG. 3.**
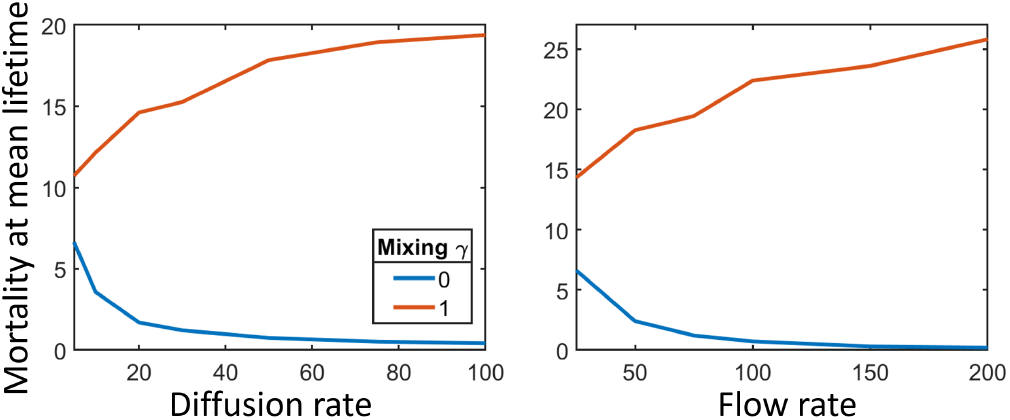
Coupled effects of diffusion and flow on purely cooperative and antagonistic systems. On the left we plot the mortality, normalized by the mean mortality, at the mean lifetime, versus diffusion rate, setting flow velocity to *v* = 0. We see that as we increase diffusion, more cells are able to couple together, thus exaggerating the aging dynamics. The cooperative system (*γ* = 1) ages more and the antagonistic system (*γ* = 0) ages less as we increase diffusion. On the right we plot the same value versus flow velocity, keeping diffusion fixed at *D*_*ϕ*_ = *D*_*ψ*_ = 5. Increasing the velocity gives the same effect as increasing diffusion. Since cells are more strongly coupled, the aging behavior for each case becomes more pronounced. The hill constants were set to *k*_*ϕ*_ = *k*_*ψ*_ = 2 for each case. Other parameters were set to *α* = 100, *λ*_*ϕ*_ = *λ*_*ψ*_ = 50, and *A*_*ϕ*_ = *A*_*ψ*_ = 10.

When we increase the diffusion rate, keeping flow at zero, we see that the effect of increased diffusion is to exaggerate the aging behavior of competitive (*γ* = 0) and cooperative (*γ* = 1) systems. Larger diffusion couples more cells together, as the secreted factors (negative or positive) diffuse out further. Hence, larger diffusion leads to a larger effect, steeper mortality curves for cooperative systems and shallower curves for competitive ones.

We see the same behavior when we increase flow. We see in Fig. 3 that the effect of increased flow (keeping diffusion constant) is to exaggerate the aging behaviour for both systems. With higher flow, the secreted factors are transported further away from the cell producing them, allowing more cells to become coupled together. As with diffusion, we again see larger aging for cooperative systems and less for competitive ones.

We therefore find that how signals and factors are transported also strongly influences the aging dynamics of an organism. With these insights, we compare the results of our model to the mortality curves of various species in the next section.

### Fits to empirical data

We next compare mortality curves obtained from our diffusive model to those from empirical data obtained from Jones, Owen R., et al. (2014) [3] in Fig. 4. As in [3], mortality curves are plotted from the age of reproductive maturity up to the terminal age where only 5% of the population is alive. Mortality values are scaled relative to their mean values for each species.

**FIG. 4.**
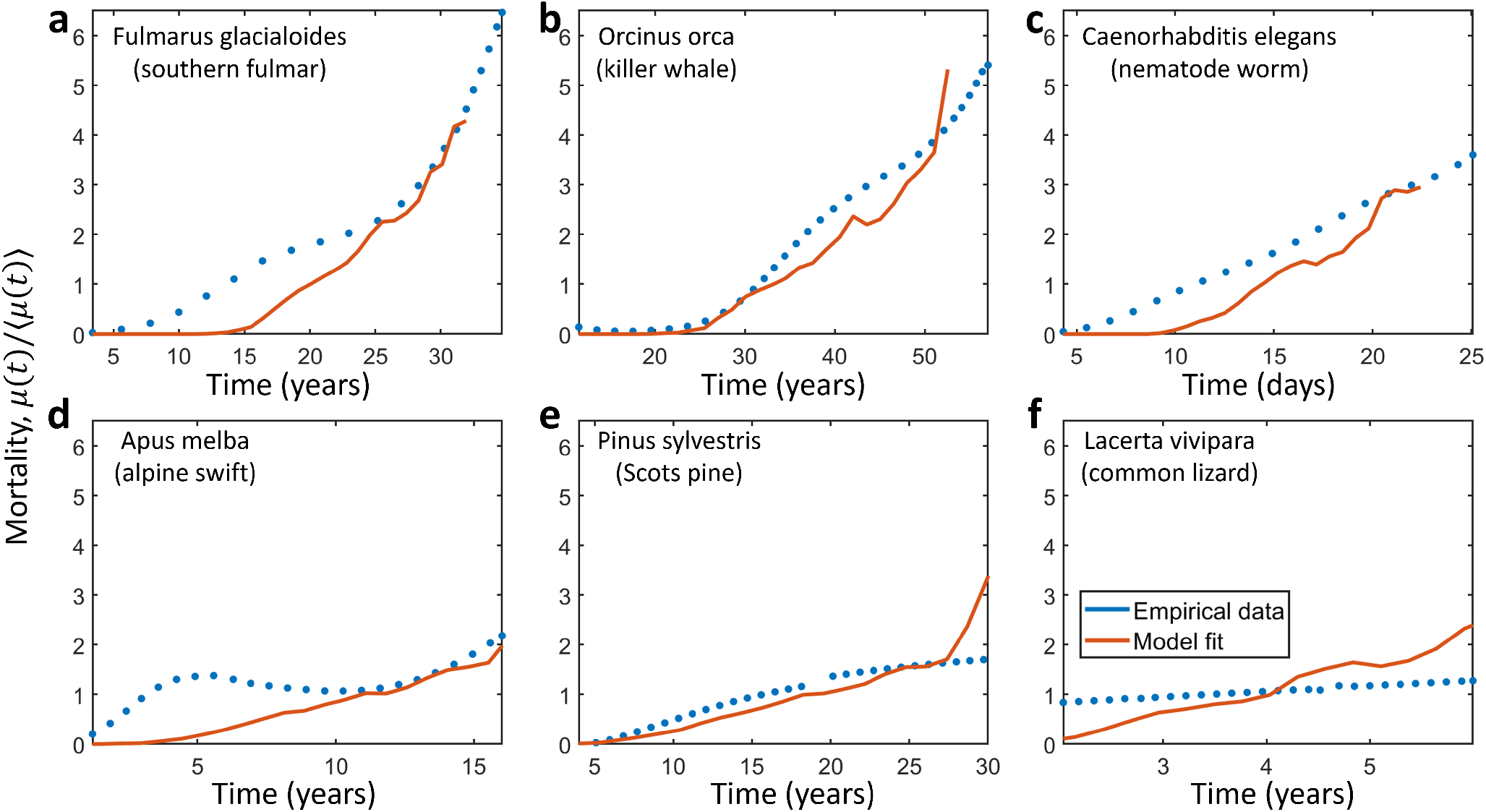
Mortality curves obtained from empirical data and fitted with theoretical model. Empirical mortality data was obtained from [3]. 7 Mortality curves are plotted as functions of age from the age of reproductive maturity up to the age where 5% of the population is still alive. Mortality values are normalized by the mean value for each case. **a**, The bird *Fulmarus glacialoides* has a steep mortality curve, with mortality at terminal age at around 6 times the average level of adult mortality. Fit parameters are *v* = 150, *k*_*ϕ*_ = 1, *k*_*ψ*_ = 1, and *γ* = 0.15. **b**, The mammal *Orcinus orca* have a slighly less steep mortality curve with the mortality at terminal age around 5 times the average level of adult mortality. Fit parameters were *v* = 150, *k*_*ϕ*_ = 2, *k*_*ψ*_ = 2, and *γ* = 0.125. **c**, *Caenorhabditis elegans*, an invertibrate, has an even less steep mortality curve, with the mortality at terminal age at around 2 times the average mortality. Fit parameters are *v* = 150, *k*_*ϕ*_ = 3, *k*_*ψ*_ = 3, and *γ* = 0.075. **d**, The bird *Apus melba*, has mortality at terminal age at around 3 times the average mortality. Fit parameters are *v* = 0, *k*_*ϕ*_ = 1, *k*_*ψ*_ = 3, and *γ* = 0.02. **e**, Mortality curve and fit for *Pinus sylvestris* (Scotts pine). Fit parameters are *v* = 0, *k*_*ϕ*_ = 1, *k*_*ψ*_ = 3, and *γ* = 0.01. **f**, Mortality curve and fit for the lizard *Lacerta vivipara*. Fit parameters are *v* = 150, *k*_*ϕ*_ = 1, *k*_*ψ*_ = 2, and *γ* = 0.0. Other parameters were set to *D*_*ϕ*_ = *D*_*ψ*_ = 5, *λ*_*ϕ*_ = *λ*_*ψ*_ = 50, and *A*_*ϕ*_ = *A*_*ψ*_ = 10. The time constant was set to *α* = 100 for all simulations and the mortality curves were rescaled to have the same time at terminal age.

Phylogenetic relatedness is seen to have some correlation with the aging dynamics of species (c.f. Fig. 1 of [3]). To compare our model to empiric data, we took a representative species from mammals (*Orcinus orca*), birds (*Fulmarus glacialoides* and *Abus melba*), invertebrates (*Caenorhabditis elegans*), plants (*Pinus sylvestris*) and reptiles (*Lacerta vivpara*) and fit mortality curves from our model in Fig. 4. We compared empiric data to our model by running simulations varying the mixing factor *γ*, flow velocity *v*, and the hill constants *k*_*ϕ*_ and *k*_*ψ*_. We then compared the resulting mortality curves and took the best fitting curve.

The bird *Fulmarus glacialoides* has a fairly steep mortality curve with a mortality rate at terminal age (33 years) at around 6 times the average mortality. Our best fit model for *Fulmarus glacialoides* is given by the values *v* = 150, *k*_*ϕ*_ = 1, *k*_*ψ*_ = 1, and *γ* = 0.15 (Fig. 4a). The mammal *Orcinus orca* has a slighly less steep mortality curve with a mortality at terminal age (59 years) at around 5 times the average. Our best fit model for *Orcinus orca* is given by the values *v* = 150, *k*_*ϕ*_ = 2, *k*_*ψ*_ = 2, and *γ* = 0.125 (Fig. 4b). The invertebrate *Caenorhabditis elegans* has a mortality at terminal age (25 days) at around 3 times the average mortality. We fit our model with values *v* = 150, *k*_*ϕ*_ = 3, *k*_*ψ*_ = 3, and *γ* = 0.075 in Fig. 4c. The bird *Apus melba* has a much smaller mortality at terminal age (16 years) compared to the fulmar, at around 2 times the average mortality. We fit our model with values *v* = 0, *k*_*ϕ*_ = 1, *k*_*ψ*_ = 3, and *γ* = 0.02 in Fig. 4d. The plant *Pinus sylvestris* has a mortality at terminal age (30 years) at around twice the average mortality. We fit our model with values *v* = 0, *k*_*ϕ*_ = 1, *k*_*ψ*_ = 3, and *γ* = 0.01 in Fig. 4e. Finally, the reptile *Lacerta vivpara* (common lizard) has a roughly flat mortality curve with mortality rate at terminal age (6 years) just slighly above the average mortality. We fit our model with values *v* = 150, *k*_*ϕ*_ = 1, *k*_*ψ*_ = 2, and *γ* = 0.0. in Fig. 4e.

In general mammals tend to have the most drastic aging behavior, reptiles tend to age less (some may even exhibit negligible or negative senescence), and plants seem to age even less. Whereas, birds and invertebrates are more diverse in their aging behavior [3]. Here we fit our model varying parameters for the mixing factor *γ*, and hill constants for each type of factor *k*_*ϕ*_ and *k*_*ψ*_. We see that we are able to find good fits to the empiric data by varying these parameters, which correspond to how strongly cells may depend on each other within the organism.

We see that species that age less, with shallower mortality curves, are fit with smaller values for the *γ* factor, corresponding to less cooperative interactions between components. Therefore, we see that the biophysical details of the interactions between cells may contribute to the diversity in aging across different species.

## DISCUSSION

It has been proposed that flatter mortality curves (such as that of reptiles and amphibians) [3] may be due to a combination of factors such as metabolism and thermal regulation, regenerative capabilities [22], fertility and life cycles [23–25], and modular structure [26], due to both evolutionary and mechanical differences across species. There have been a few theoretical studies aiming to explain the diversity of aging. Indeterminate growth has been proposed as a factor leading to constant or negative senescence. If mortality decreases with increasing size and reproductive potential rises, non-senescence can become optimal evolutionarily [2]. Indeterminate growth may then explain why plants and some reptiles seem to age slower than species with determinate growth.

Here we consider an additional, possibly alternative mechanism behind the phylogenetic trends in aging: We hypothesize that how various factors are transported between cells determines the nature of *µ*(*t*).

We studied two separate models to explore the effects of interdependencies on aging. In the first model, we emphasized the connectivity structure, and study aging processes on ensembles of different network topologies. We found in general that sparser, locally connected topologies give rise to shallower mortality curves, whereas denser, globally connected networks give steeper mortality curves. We also found the initial conditions more largely affect denser networks. Poor initial conditions greatly increase the probability of failure for denser networks, with little room to increase over time. This then leads to shallower mortality curves (Fig. 1e-h). For sparser networks, we find a larger repair rate can lead to shallower mortality curves (Fig. 1c,d), as opposed to denser networks where the repair rate has little effect. As failure propagates slower in sparser networks, they are more amenable to repair.

In the second model, we emphasized the biophysical properties of the signals being communicated. In this diffusion-advection model, cells influence each other’s function through secreted factors that are able to diffuse and flow. In contrast to network models, we studied a simple circular topology, but were able to focus on the inter-cellular biophysical processes in detail. Even with this simple topology, we find that differences in the types of factors and how those factors propagate can strongly determine the mortality curves of a species.

We assumed cells communicate through two types of secreted factors, a cooperative factor *ϕ* such as antioxidant enzymes or growth or survival factors [18–21], and an antagonistic factor *ψ* such as waste and apoptotic factors [27, 28]. We modeled the relative ratio of factors through the mixing parameter *γ*, with *γ* = 1 corresponding to a purely cooperative system and *γ* = 0 a purely antagonistic one.

We found the aging behavior of the system is largely determined by the factor *γ* as well as by the hill constants *k*_*ϕ*_ and *k*_*ψ*_. Lower values of *γ*, corresponding to less cooperative interactions between cells in a species, give shallower mortality curves, and higher values of *γ* give steeper mortality curves (Fig. 2). This is because for larger *γ*, cells depend on each other for survival. As damages accumulate, a late life “crash” becomes more likely, where all cells die out, killing the organism. For smaller *γ*, cells cooperate less and may even compete with each other. Accumulated damages then do not affect other cells as much and system wide death is less drastic. Larger values for the hill constants further exaggerate these effects, giving a larger difference between small and large values of *γ* when the hill constants are also larger (Fig. 2).

We also explored the effects of diffusion and flow on aging. We found larger diffusion and flow also exaggerate aging effects (Fig. 3). A larger diffusion and/or flow rate couples more distant cells together. Cells then cooperate more for large *γ* or compete more for small *γ*, leading to more steep or shallow mortality curves, respectively.

Finally, we compared the results of our theoretical model by fitting mortality curves to empirical data obtained from Jones, Owen R., et al. (2014) [3]. We plotted empirical mortality curves and theoretical fits for mammals (*Orcinus orca*), birds (*Fulmarus glacialoides* and *Apus melba*), invertebrates (*Caenorhabditis elegans*), plants (*Pinus sylvestris*), and reptiles (*Lacerta vivpara*) in Fig. 4. Fit values for *γ* corresponded to larger *γ* for species with steeper mortality curves and smaller *γ* for shallower ones, as expected. We therefore find that the biophysical details of how various factors are transported within an organism can strongly determine its aging dynamics.

In reality, there may be many diffusive factors that determine the aging dynamics of an organism. These may also be integrated in various ways beyond what we consider here. We expect that multiple factors with similar diffusion-advection length scales would not significantly change our results, and the factors *ϕ* and *ψ* considered here can be taken as a sum of such factors or as limiting factors. Exploring different ways of integrating factors may give new and interesting results beyond what we consider here.

We also took in our model, for simplicity, each type of factor to be produced by all cells at a constant rate. In reality, the production of factors may depend on the physiolgoical state of the cell or on some feedback mechanism. Factors produced by cell death or lysis might also be relevant to include. Local, non-diffusive compounds such as lipofuscin [29] may also contribute to aging [30]. The inclusion of these additional types of factors might lead to interesting future research.

For further investigation, we take can also take into account cellular reproduction and repair in the diffusion-advection model. This may produce flatter and even negatively aging mortality curves as seen in some plants and reptiles. It would also be interesting to see the effect cell health has on flow. For example, the integrity and tension of the extra-cellular matrix which helps to regulate the interstitial fluid pressure depends on cellular viability [31]. It may be insightful to also include the mechanical interplay of flow and cell viability in a more complete theory of aging.

Finally, it could be interesting to combine the two models explored here, into a biophysically realistic network approach, extending our current one-dimensional diffusion-advection model to more realistic topologies. This approach may also give further insight into the aging dynamics of various species.

